# Validating Neurite EXchange Imaging (NEXI) using diffusion Monte Carlo simulations in realistic numerical gray matter substrates

**DOI:** 10.64898/2026.02.11.705314

**Authors:** Rita Oliveira, Jasmine Nguyen-Duc, Malte Brammerloh, Ileana Jelescu

## Abstract

NEXI is a gray matter (GM) microstructural model designed to probe brain tissue microstructure *in vivo* using diffusion MRI. NEXI describes GM as two exchanging Gaussian compartments – neurites, modeled as randomly oriented, infinitely long sticks, and the extracellular space – allowing the estimation of biophysically interpretable parameters related to neurite microstructure and intercompartmental exchange. While modeling cell processes as sticks and each compartment as Gaussian are common assumptions for brain biophysical models of diffusion, neurite structural irregularities and the presence of somas, particularly in GM, may violate them and bias NEXI parameter estimates. Furthermore, the barrier-limited exchange assumed in the Kärger model that underlies NEXI may also be violated in realistic conditions. Therefore, in this work, we evaluate NEXI’s accuracy in numerical substrates that incorporate realistic GM features and membrane permeability. To this end, we generated several GM-like substrates with neurite beading, undulation, orientation dispersion, and somas across a range of membrane permeabilities. Diffusion signals were generated with Monte Carlo simulations of water diffusion and subsequently fitted with NEXI. Overall, NEXI accurately recovered exchange times across permeability levels and successfully disentangled exchange effects from other microstructural features, showing only minor bias in estimates from the realistic geometries. These results support its potential for *in vivo* GM microstructure mapping and studies of brain disorders.

## 1 Introduction

Characterization of brain tissue microstructure *in vivo* from Magnetic Resonance Imaging (MRI) has long been a major focus in neuroscience, as it enables non-invasive investigation of the structural underpinnings of brain function in health and disease (Novikov et al., 2018; Weiskopf et al., 2015, 2021). Diffusion MRI, in particular, is a powerful tool for this purpose, as the signal reflects the motion of water molecules and the constraints imposed by the underlying tissue architecture. However, because diffusion MRI acquisitions have a millimeter-scale resolution, they cannot directly resolve microscopic features of brain tissue, such as cell size. To bridge this gap, biophysical models link the diffusion MRI signal to parameters that describe the underlying tissue in a simplified geometry. This non-invasive approach, often referred to as *Microstructural MRI* (Jelescu et al., 2020; Novikov et al., 2018; Weiskopf et al., 2015), has become an important framework for probing brain tissue microstructure *in vivo*.

Within this framework, several biophysical models have been proposed, each relying on different assumptions and tailored to specific tissue types. In white matter (WM), models such as the Standard Model (SM) (Coelho et al., 2024; Novikov et al., 2018) have been widely adopted. In gray matter (GM), commonly used models include the Soma and Neurite Density Imaging (SANDI), which extends the SM by adding a spherical compartment to represent cell bodies (Palombo et al., 2020), and Neurite Exchange Imaging (NEXI) (Jelescu et al., 2022) or Standard Model with Exchange (SMEX) (Olesen et al., 2022), which includes water exchange between cell processes (dendrites, axons, and glial processes) and the extracellular space.

More specifically, the NEXI/SMEX (Jelescu et al., 2022; Olesen et al., 2022; Uhl et al., 2024) model of GM has two exchanging compartments: a collection of neurites modeled as randomly oriented infinitely long sticks, and the extracellular space modeled as an isotropic Gaussian medium. They are based on Kärger’s analytical framework of barrier-limited exchange between two Gaussian compartments. They differ in that the NEXI signal equations were derived under the narrow pulse approximation, whereas SMEX explicitly accounts for the finite width of the diffusion gradients. Fitting NEXI/SMEX to diffusion MRI data yields estimates of neurite water fraction (*f*), intra- and extracellular diffusivities (*D_i_* and *D_e_*, respectively), and exchange time between the neurite and extracellular spaces (*t_ex_*), all of which are of potential clinical or biological relevance. In particular, the exchange time is assumed to relate to cell membrane permeability, and has been shown to correlate well (but not exclusively) with myelination in the cortex (Hertanu et al., 2025; Uhl, Pavan, Feiweier, et al., 2025).

NEXI has been successfully applied in rat brains *in vivo* (Jelescu et al., 2022) and *ex vivo* (Olesen et al., 2022), and human brains both *post mortem* (Hertanu et al., 2023) and *in vivo*, using the Connectome (Chan et al., 2025; Uhl, Pavan, Gerold, et al., 2025; Uhl et al., 2024) and clinical scanners (Hertanu et al., 2025; Uhl, Pavan, Feiweier, et al., 2025). These studies demonstrate the feasibility of estimating human cortical microstructural parameters *in vivo*.

Despite NEXI’s demonstrated potential, the validity of its assumptions remains insufficiently characterized. Structural features commonly observed in brain tissue, such as neurite beading and undulation, may violate the Gaussian diffusion assumption of each compartment underlying the Kärger model (H.-H. Lee et al., 2020; Nguyen-Duc, Brammerloh, et al., 2026; Novikov et al., 2014), thereby affecting the accuracy of the estimated NEXI microstructural parameters. Additional cellular features, such as somas (Olesen et al., 2022) and dendritic spines (Chakwizira et al., 2025; Şimşek et al., 2025), can further influence the accuracy of NEXI-derived microstructural estimates. In particular, exchange between dendritic spines and the shaft have recently been shown to produce an MRI signal signature similar to that due to inter-compartmental exchange across the membrane, although ultra-strong diffusion gradients (preclinical-grade and higher) would be needed to observe this effect experimentally (Chakwizira et al., 2025; Şimşek et al., 2025).

Previous validation efforts for NEXI have not accounted for these effects: some studies generated synthetic data and fitted it using the same NEXI analytical model (Chan et al., 2025; Jelescu et al., 2022; Uhl et al., 2024), while others employed numerical phantoms with realistic neuronal geometries but restricted the MRI signal to the intracellular compartment, thereby excluding exchange with the extracellular space (Olesen et al., 2022; Şimşek et al., 2025).

To address such limitations, numerical phantoms of packed cells into a synthetic tissue voxel have emerged as a powerful framework for validating biophysical models (Fieremans & Lee, 2018; Ginsburger et al., 2019; Nguyen-Duc, Brammerloh, et al., 2026). They allow the generation of tissue-like substrates with known ground-truth properties, in which the diffusion process can be simulated, typically with Monte Carlo methods. The resulting synthetic MRI signals can then be fitted with biophysical models, allowing a direct comparison between estimated parameters and their true ground-truth values (Jelescu et al., 2020; Nguyen-Duc, Uhl, et al., 2026). Importantly, numerical phantoms are particularly useful for studying permeability, as water exchange across the cellular membrane can be explicitly modeled and systematically varied (Gardier et al., 2023).

In this work, we use numerical phantoms to evaluate the performance of NEXI in GM-like substrates packed with beaded and undulating neurites, and spanning a range of membrane permeabilities. We additionally included somas, which are not accounted for in the NEXI model, and varied the level of neurite orientation dispersion to reflect different GM architectures. We did not include dendritic spines in our simulations, as previous work has shown that spine-related sensitivity arises primarily at short diffusion times (Şimşek et al., 2025), which are generally not accessible with the clinical or preclinical diffusion MRI sequences used for NEXI. In addition, reliable numerical simulators explicitly incorporating spines are currently not available.

## 2 Methods

### 2.1 Numerical substrate generation

Numerical substrates representing GM were generated within a (100 *µ*m)^3^ voxel using the CATERPillar toolbox (Nguyen-Duc, Brammerloh, et al., 2026) (v. 05/08/2025). This generator models neurites as chains of overlapping spheres.

The generated substrates consisted of three orthogonal fiber populations to reproduce the high orientation dispersion expected in GM (Lampinen et al., 2020; Salo et al., 2021), with a target neurite volume fraction (*f*) of 65%. The neurite radius distribution was aimed to peak around 1 *µ*m, in agreement with reported values of 0.5–1 *µ*m in humans and rodents (Aird-Rossiter et al., 2026; Palombo et al., 2019). Since CATERPillar inflates neurites in a final step to achieve the target volume fraction, the substrate generation was initialized with a gamma distribution of radii using a shape parameter *α* = 1.60 and a scale parameter *β* = 0.30 (corresponding to a mode of approximately 0.18 *µ*m) for all substrates.

To increase the realism of neurites, we incorporated two morphological features: beading (local radius fluctuations along the neurite) and undulation (deviation of the centerline from a straight path).

In CATERpillar, beading amplitude *A* is defined as a fraction of the neurite’s initial radius. In this work, neurite beading is quantified by the coefficient of variation of the radius 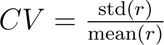(H. H. Lee et al., 2019; H.-H. Lee et al., 2024). As a representative baseline value, we set CATERPillar parameter *A* to 0.2 times the radius, which corresponds to a CV of ∼ 0.2. This choice is motivated by electron microscopy studies of axons (as quantitative data on neurite beading are scarce), which report beading amplitudes of 0.2–0.3 times the radius (CV∼ 0.2 (H. H. Lee et al., 2019; H.-H. Lee et al., 2024)), with substantially larger beading amplitudes of up to ∼ 0.57 times the radius observed only under pathological conditions such as ischemia (Budde & Frank, 2010).

In CATERPillar, undulation is controlled by parameter *ε*, which modulates the strength of attraction toward a given target (Nguyen-Duc, Brammerloh, et al., 2026). In this work, we use the term ‘undulation’ (und) to refer to the ratio of total process length divided by its end-to-end distance, rather than tortuosity – a commonly used term for processes undulation (Aird-Rossiter et al., 2026) – to avoid confusion with extracellular tortuosity which is defined as the ratio of free diffusivity to effective hindered diffusivity (e.g. *D_free_/D_e_*) (Jespersen et al., 2007; Kaden et al., 2016; H.-H. Lee et al., 2020; Novikov, 2021). As a representative baseline value, we set CATERPillar *ε* to 0.2, which yields an undulation factor of ∼ 1.10. This value is consistent with microscopy-reconstructed neurons, which report median branch undulation factors of ∼ 1.30 −1.35 for for pyramidal and GABAergic neurons in humans (Aird-Rossiter et al., 2026), and 1.10–1.15 for dendrites and 1.10–1.20 for axons in rats (Stepanyants et al., 2004).

This substrate was considered the most realistic and used as the default (Table 1). It was then systematically modified by varying individual CATERPillar parameters: beading amplitude *A* from 0 to 0.5 times the initial radius (step 0.1), corresponding to CV=0.18-0.34; undulation amplitude *ε* from 0 to 0.4 (step 0.1), corresponding to undulation factors 1.00-1.30 (Table 1). The level of neurite orientation dispersion is characterized by the *l* = 2 rotational invariant *p*_2_ and related to the dispersion angle Ψ by 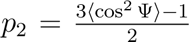 (with *p*_2_ = 0 corresponding to a fully isotropic orientation distribution and *p*_2_ = 1 to perfectly aligned neurites). Orientation dispersion was modified by reducing the number of orthogonal fiber populations from three to two. Because neurite growth and packing is easier under certain morphological conditions, requiring less inflation in the final step to maintain a comparable radius distribution and volume fraction, we used shape and scale parameters of *α* = 5.00 and *β* = 0.20 for the radius distribution, with a neurite volume fraction of 40% for the non-undulated case, and *α* = 4.00 and *β* = 0.20 for the two-fiber case.

**Table 1:** Target CATERPillar substrate parameters, including beading (amplitude *A* and expected final CV), undulation (*ɛ* and expected final undulation ratio *und*), number of fiber populations, soma fraction *f_s_*, and neurite membrane permeability *κ* (*µ*m*/*s).

| Condition | Beading | | Undulation | | # Fibers | $f_s$ | $\kappa$ ($\mu\text{m/s}$) |
| --- | --- | --- | --- | --- | --- | --- | --- |
| | $A$ | CV | $\epsilon$ | $und$ | | | |
| Default | 0.2 | $\approx 0.24$ | 0.2 | $\approx 1.10$ | 3 | 0 | 20 |
| Permeability increase | 0.2 | $\approx 0.24$ | 0.2 | $\approx 1.10$ | 3 | 0 | 0–30 |
| Beading increase | 0–0.5 | $\approx 0.18$–$0.34$ | 0.2 | $\approx 1.10$ | 3 | 0 | 20 |
| Undulation increase | 0.2 | $\approx 0.24$ | 0–0.4 | $\approx 1.0$–$1.30$ | 3 | 0 | 20 |
| Orientation dispersion decrease | 0.2 | $\approx 0.24$ | 0.2 | $\approx 1.10$ | 2–3 | 0 | 20 |
| With somas | 0.2 | $\approx 0.24$ | 0.2 | $\approx 1.10$ | 3 | 0.05–0.1 | 20 |

For the default substrate, we evaluated several levels of membrane permeability: *κ* = 0, 10, 20, and 30 *µ*m*/*s, in agreement with experimental membrane permeability values in healthy cells (Boss et al., 2013; Harkins et al., 2009; Stanisz et al., 1997). For the remaining substrates, *κ* was kept at 20 *µ*m*/*s. Finally, we also generated two substrates that included somas (Table 1). These substrates shared the same features as the default configuration but incorporated a population of spherical somas uniformly distributed in the voxel. The soma volume fractions (*f_s_*) in these two substrates were 5% and 10%, respectively, and the soma radius distribution had mean 7.1 *µ*m and standard deviation 3.6 *µ*m, consistent with reported values for pyramidal cells in rats (Aird-Rossiter et al., 2026; Olesen et al., 2022). In our substrates, somas were disconnected from neurites, as the former were placed using CATERPillar’s glial cell population generator and the latter using the axon population generator (as CATERPillar is primarily designed for WM substrates). Nonetheless, having disconnected somas also enables us to ensure comparability of this particular substrate with the others, as the main difference is indeed only the addition of spherical components. Furthermore, soma-dendrite exchange is typically quite slow (De Riedmatten et al., 2024). In these substrates, soma membranes were first assumed impermeable, as in SANDI (Palombo et al., 2020), and then also assigned a permeability of 20 *µ*m*/*s, the same as the neurites’.

A summary of the most important parameters used to generate the substrates is presented in Table 1.

### 2.2 Monte Carlo Simulations

To estimate MRI signals from the generated synthetic substrates, we performed Monte Carlo simulations of water diffusion. We used a simulator adapted to permeable cell membranes between intra- and extracellular spaces (Gardier et al., 2023; Rafael-Patino et al., 2020). The simulator was further modified to handle cellular structures defined by overlapping spheres instead of triangular meshes, as implemented for CATERPillar (Nguyen-Duc, Brammerloh, et al., 2026; Nguyen-Duc, Uhl, et al., 2026).

The trajectory simulations were performed for a total time of *TE* = 52 ms. The intrinsic diffusivities in the intra- (*D_i_*) and extracellular (*D_e_*) spaces were set to 2 *µ*m^2^*/*ms and 1 *µ*m^2^*/*ms, respectively, in line with the expected diffusivities (Jallais et al., 2025; Jelescu et al., 2022; Uhl, Pavan, Feiweier, et al., 2025) and supporting the condition *D_i_ > D_e_* (Dhital et al., 2019; Jelescu et al., 2020; Kunz et al., 2018). Each simulation was initialized with a walker density of 1 walker */µm*^3^, corresponding to 10^6^ walkers, distributed homogeneously inside and outside the neurites. This particle density was in line with previously reported benchmark values (Fieremans & Lee, 2018) and was further confirmed by the stability of the estimated metrics with increasing walker density (Fig. S1). The step size of each walker in one direction *δx* was set to ~ 0.06 *µ*m, approximately one-tenth of the minimum neurite radius on the final substrates. The corresponding simulation step duration Δ*t* was = 1 *µ*s, computed from 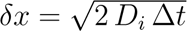, leading to a total number of steps *T* = 64800 (as Δ*t* = *TE/T*).

Synthetic diffusion-weighted signals were generated from the resulting trajectory simulations assuming a pulsed-gradient spin-echo (PGSE) sequence and a realistic preclinical (*G_max_* ∼ 600 mT/m) *in vivo* protocol (Jelescu et al., 2022), with *b*-values of 1 (12 directions), 2 (16 directions), 3.5 (24 directions), 5 (30 directions), and 7 ms*/µ*m^2^ (40 directions). The diffusion times (Δ) were 15, 26, and 38 ms, with a gradient duration (*δ*) of 4.5 ms to be within the narrow pulse approximation. The echo time was *TE* = 52 ms (but no *T*_2_ relaxation was modeled).

We further evaluated NEXI performance under a clinically feasible protocol (characterized by limited gradient performance, which constrains the achievable b-values and requires longer *δ* and Δ). Synthetic diffusion-weighted signals were generated using a clinical diffusion protocol following Uhl, Pavan, Feiweier, et al., 2025. Specifically, diffusion times (Δ) of 28.3, 36, 45, 55, and 65 ms were simulated. For Δ = 28.3 ms and 36 ms, *b*-values of 1 and 2 ms*/µ*m^2^ were used; for Δ = 45 ms, *b*-values of 1, 2, 3.2, and 4.44 ms*/µ*m^2^; and for Δ = 55 ms and 65 ms, *b*-values of 1, 2, 3.2, and 5 ms*/µ*m^2^. Each Δ included 20 gradient directions per shell, with *δ*=16.5 ms. In the Monte Carlo simulations, the number of steps *T* was adjusted such that *TE* = 100 ms.

### 2.3 Analysis

The NEXI model was fitted to the magnitude of the generated complex-valued diffusion-weighted signals using the implementation available in the Swiss Knife Toolbox (https://github.com/QuentinUhl/ graymatter swissknife (Jelescu et al., 2022; Uhl, Pavan, Feiweier, et al., 2025; Uhl et al., 2024)). In the case of impermeable substrates (*κ* = 0), *t_ex_* was also fixed to an arbitrarily high value in the fitting procedure reflecting the absence of inter-compartmental exchange, while the remaining parameters were estimated. NEXI estimates for *f*, *D_i_*, *D_e_* and *t_ex_* were compared to ground-truth values of the substrate. Ground-truth values for *f* were directly obtained from the geometric properties of the neurites in the substrate; for *t_ex_*, they were derived from the relationship between membrane permeability and neurite surface-to-volume ratio (*S/V*), given by 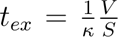 (Fieremans & Lee, 2018; H.-H. Lee et al., 2025). Ground-truth values for *D_i_* and *D_e_* were computed from the walker trajectories obtained for each substrate without permeability.

To further characterize the microstructure underlying the simulated voxel, we assessed the time dependence of mean diffusivity (MD) and mean kurtosis (MK). To this end, diffusion kurtosis imaging (DKI) was performed independently at each diffusion time using the TMI toolbox (Veraart et al., 2013).

For comparison with the full NEXI model estimate, exchange time *t_ex_* was also computed from the MK time dependence (Fieremans et al., 2010; Jensen & Helpern, 2010):

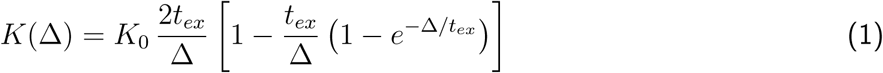

To evaluate estimation errors under realistic noise conditions, we added Rician noise by introducing Gaussian noise to the real and imaginary parts of the noise-free signals and then taking the magnitude. This was done for signals generated at different permeability levels in the default substrate, with an SNR of 50. The procedure was repeated 500 times, and the data from each realization were fitted as described above. In these cases, in the NEXI fit, the Rician noise floor was accounted for (Uhl et al., 2024).

In the case of the clinical protocol, SMEX, which accounts for the non-zero duration of diffusion gradient pulses, was fitted instead of NEXI.

## 3 Results

### 3.1 Numerical substrates generation

The generated substrates broadly matched the targeted morphological parameters (Fig. 1), with a few expected deviations. Final neurite radii were slightly larger than the target distribution due to the inflation step applied in CATERPillar to reach the desired volume fraction. By undershooting the target distribution, we obtained a final mean radius within the reported range of 0.5–1 *µ*m in humans and rodents (Aird-Rossiter et al., 2026; Palombo et al., 2019). The target neurite volume fraction of 75%, however, was not achieved, with the final value ranging 31 - 53%, depending on the substrate (Fig. 1). This limitation arises from the difficulty of densely packing neurites (a challenge commonly encountered in GM substrate generation (Fieremans & Lee, 2018; Ginsburger et al., 2019)), particularly when incorporating three different fiber orientations, as in our case. Nevertheless, the achieved neurite volume fractions across substrates are comparable to literature neurite density estimates *in vivo* using either NEXI or SANDI (Jelescu et al., 2022; Schiavi et al., 2023; Uhl, Pavan, Feiweier, et al., 2025).

**Figure 1:**
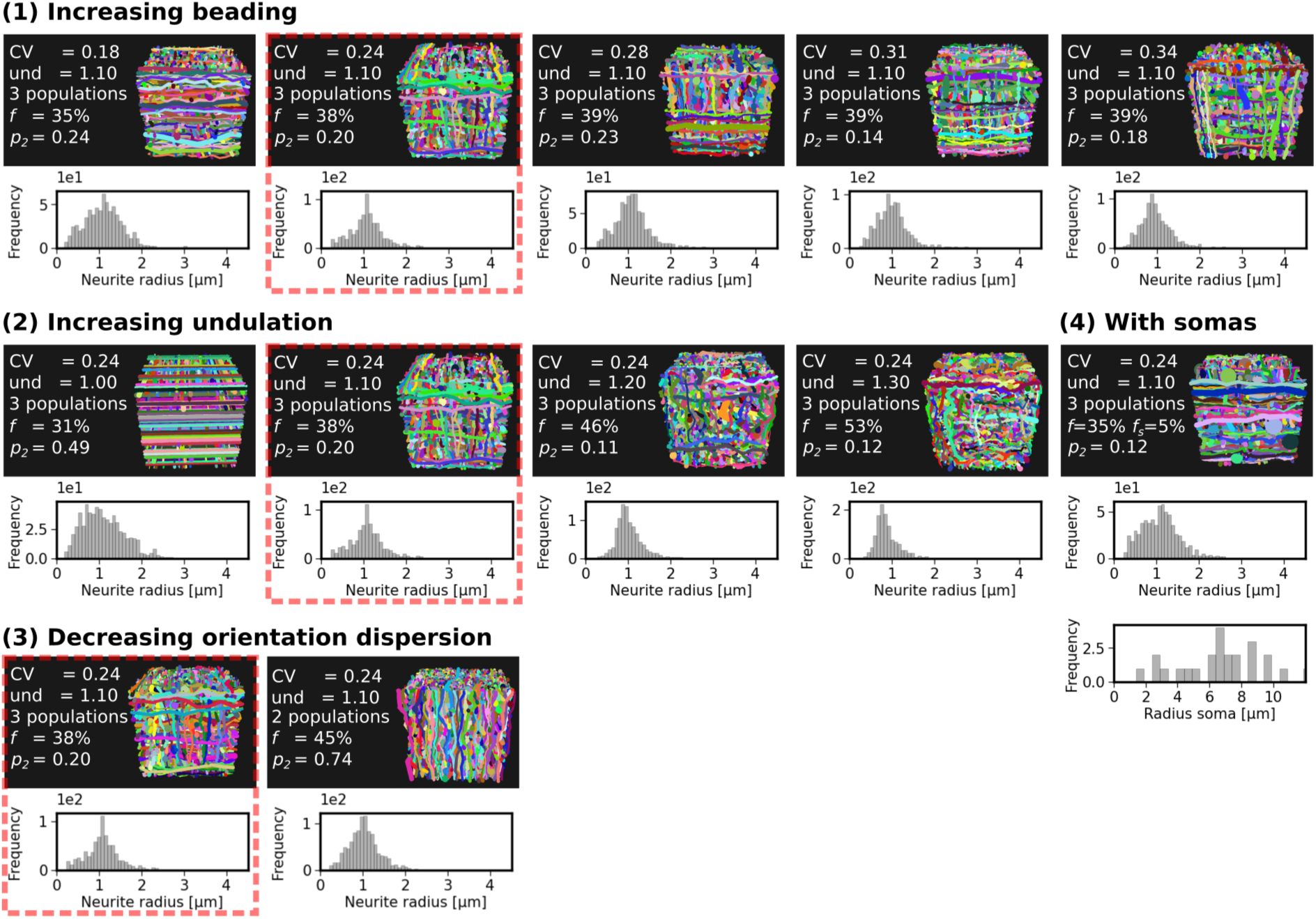
Generated substrates. Substrates mimicking GM were generated with different morphological features: (1) increasing beading level, (2) increasing undulation level, (3) decreasing orientation dispersion, and (4) presence of somas. The default substrate is highlighted in red. Beading level was defined as the coefficient of variation of the radius, given by 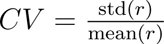 (H. H. Lee et al., 2019), and undulation was defined as the ratio between total neurite length and its end-to-end distance; both averaged across neurites. The resulting neurite volume fraction (*f*), soma volume fraction (*f_s_*), orientation dispersion (*p*_2_), the distribution of mean radius along each neurite, and distribution of soma radius are shown for each case.

The different substrates were broadly comparable, particularly in terms of the neurite radius distribution. Orientation dispersion, quantified by *p*_2_, fell within the expected range for GM (∼ 0.13–0.30) (Lampinen et al., 2020; Salo et al., 2021) for most substrates. Removing undulation increased *p*_2_, as neurites became straighter and exhibited less local directional variability. The neurite volume fraction *f* increased slightly with stronger beading (reflecting the larger neurite volume) and with larger undulation (as wavier neurites take up more space and can grow around obstacles, facilitating denser packing). Note that even when beading is not introduced (*A* = 0), neurites still undergo local constrictions to pass through narrow gaps, leading to a final CV different from 0 (= 0.18).

### 3.2 Impact of cell membrane permeability

First, we evaluated NEXI performance on the default substrate (beading CV = 0.24, undulation = 1.10) across varying membrane permeabilities.

The estimated *t_ex_* ranged from 15 to 49 ms and showed close agreement with theoretical predictions (Fig. 2a), with biases of -13%, -20%, and -21% for permeabilities of *κ* = 10, 20, and 30 *µ*m*/*s, respectively.

**Figure 2:**
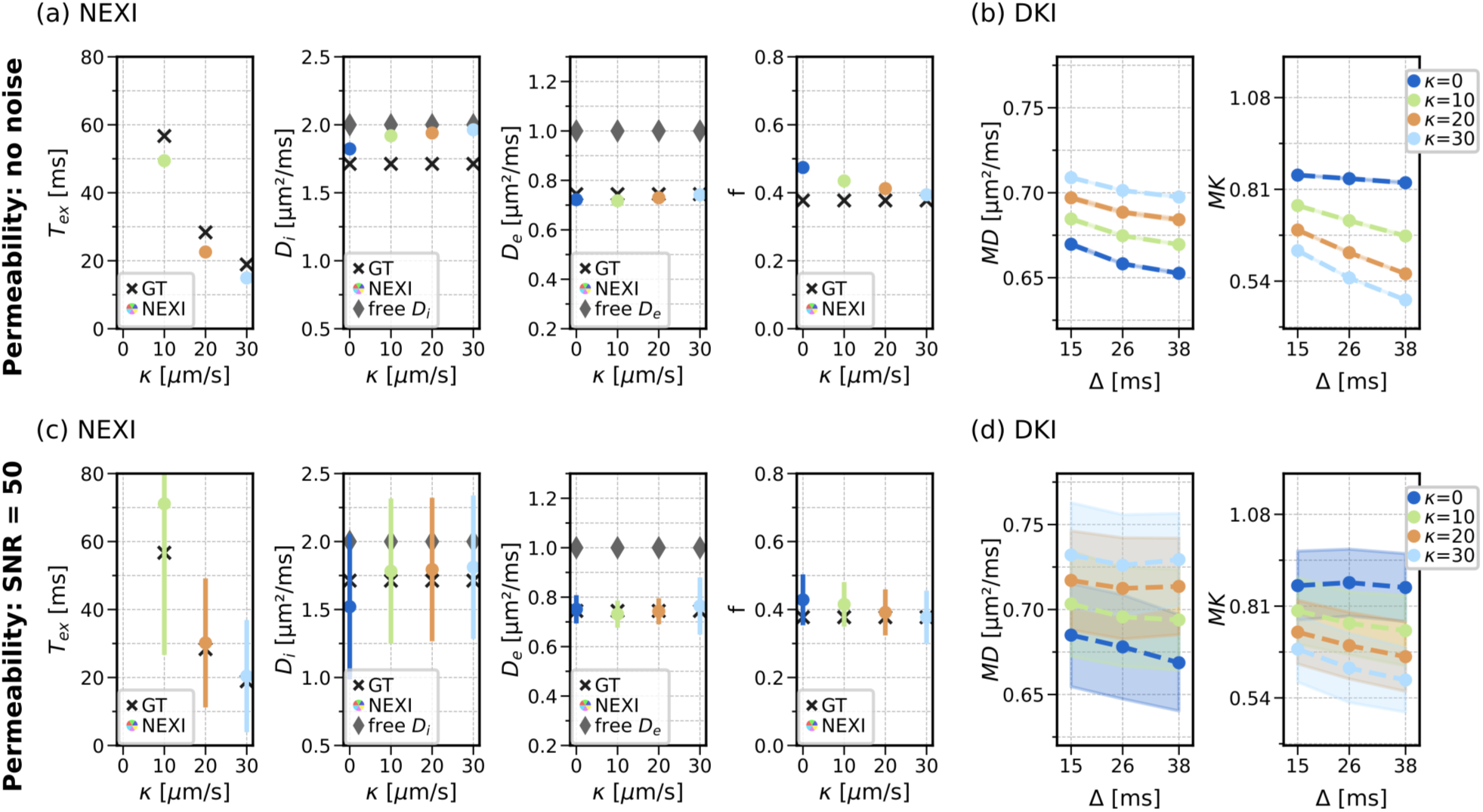
NEXI performance across permeability levels. NEXI parameters *D_i_* and *D_e_* agreed with trajectory-derived values (black markers) and were lower than the intrinsic diffusivity (gray markers), reflecting hindered diffusion due to neurite irregularities (a). *t_ex_* and *f* followed theoretical predictions, with slight deviations at low permeability (a). MD showed weak time dependence, while MK exhibited a pronounced time dependence that increased with permeability (b). With added noise, the same overall trends were preserved, but variability increased in both NEXI (c) and DKI (d) estimates. Black cross markers indicate ground-truth (GT) values and gray diamond markers indicate *D_i_* and *D_e_*, the intrinsic free diffusivities specified in the simulator.

NEXI estimates of *D_i_* and *D_e_* were lower than the intrinsic diffusivities specified in the simulator (gray markers) (Fig. 2a), as expected due to neurite irregularities and orientation dispersion, which introduce additional obstacles in both intra- and extracellular compartments and thereby lead to more hindered diffusion. *D_e_* closely matched the diffusivities computed directly from the simulated extracellular walker trajectories (black markers) and remained stable across permeability levels. In contrast, *D_i_* agreed with the intracellular trajectory-derived diffusivities only at low permeabilities and increased toward the intrinsic free diffusivity as membrane permeability increased. The estimation of *f* slightly improved with increasing permeability.

Consistent with the NEXI parameters, the diffusion kurtosis metrics also reflected the effect of exchange (Fig. 2b): MK exhibited a pronounced time dependence that increased with increasing permeability, whereas MD showed only a weak time dependence with slopes largely preserved across permeability levels (Table 2a). This behavior is in line with the expected dominant influence of exchange over structural disorder (Fieremans et al., 2010; Jelescu et al., 2022). Exchange times derived from MK time dependence were on average about 9 ms longer than the corresponding NEXI estimates (Table 3a).

**Table 2:** Slopes of MD and MK time dependence (in 10^3^ *µ*m^2^*/*ms^2^) for different permeability levels. For the case with noise, values are reported as mean standard error of the mean across noise realizations. Statistical significance was assessed using a one-sample t-test against zero; slopes significantly different from zero (p *<* 0.05) are marked with (*).

| (a) Noise free (see Fig. 2b) |  |  | (b) With noise (SNR=50) (see Fig. 2d) |  |  |
| --- | --- | --- | --- | --- | --- |
|  | slope MD | slope MK |  | slope MD | slope MK |
| $\kappa = 0 \mu\text{m/s}$ | -0.74 | -0.97 | $\kappa = 0 \mu\text{m/s}$ | $-0.86 \pm 0.16^*$ | $-0.84 \pm 0.59$ |
| $\kappa = 10 \mu\text{m/s}$ | -0.65 | -3.89 | $\kappa = 10 \mu\text{m/s}$ | $-0.67 \pm 0.17^*$ | $-3.21 \pm 0.68^*$ |
| $\kappa = 20 \mu\text{m/s}$ | -0.56 | -5.59 | $\kappa = 20 \mu\text{m/s}$ | $-0.19 \pm 0.22$ | $-3.48 \pm 0.88^*$ |
| $\kappa = 30 \mu\text{m/s}$ | -0.49 | -6.30 | $\kappa = 30 \mu\text{m/s}$ | $-0.54 \pm 0.30$ | $-1.36 \pm 1.07$ |

**Table 3:** Exchange time *t_ex_* (ms): ground-truth (GT), NEXI-estimated, and MK time-dependence–derived values for different permeability levels. For the case with noise, values are reported as mean standard error of the mean across noise realizations.

| (a) Noise free |  |  |  | (b) With noise (SNR=50) |  |  |  |
| --- | --- | --- | --- | --- | --- | --- | --- |
| | GT $t_{ex}$ | NEXI $\hat{t}_{ex}$ | MK $\hat{t}_{ex}$ | | GT $t_{ex}$ | NEXI $\hat{t}_{ex}$ | MK $\hat{t}_{ex}$ |
| $\kappa = 0 \mu\text{m/s}$ | - | - | - | $\kappa = 0 \mu\text{m/s}$ | - | - | - |
| $\kappa = 10 \mu\text{m/s}$ | 57 | 49 | 57 | $\kappa = 10 \mu\text{m/s}$ | 57 | 71±2 | 92±6 |
| $\kappa = 20 \mu\text{m/s}$ | 28 | 23 | 32 | $\kappa = 20 \mu\text{m/s}$ | 28 | 30±1 | 75±8 |
| $\kappa = 30 \mu\text{m/s}$ | 19 | 15 | 24 | $\kappa = 30 \mu\text{m/s}$ | 19 | 20±1 | 98±10 |

In a more realistic scenario with noise (SNR = 50), NEXI estimates of *t_ex_* showed bias of +44%, +34%, and +37% relative to the noise-free case for permeabilities of *κ* = 10, 20, and 30 *µ*m*/*s, respectively (Fig. 2c). Among the four parameters, *D_i_* and *t_ex_* showed the largest variability, as expected given their higher estimation uncertainty (Jallais et al., 2025; Uhl et al., 2024), and their bias was in the opposite direction compared with the noise-free scenario. Nevertheless, the ground-truth parameter values still fall within the corresponding uncertainty ranges of the estimates. Similarly, MD and MK also exhibited increased uncertainty under noisy conditions (Fig. 2d), but the expected trend of stronger MK time dependence with higher permeability was still observed (Table 2b). As expected, with a lower SNR (=20), both the variability and the bias of the NEXI estimates increased further (Fig. S2c). The time dependence of MD and MK becomes unstable and departs from the expected physical behavior (Fig. S2d). The exchange times estimated from MK time dependence deviated more from the ground-truth than the NEXI estimates (Table 3b).

### 3.3 Impact of neurite irregularities

We next evaluated the impact of neurite beading and undulation on NEXI estimates, using a reference permeability of *κ* = 20 *µ*m*/*s (Fig. 3).

**Figure 3:**
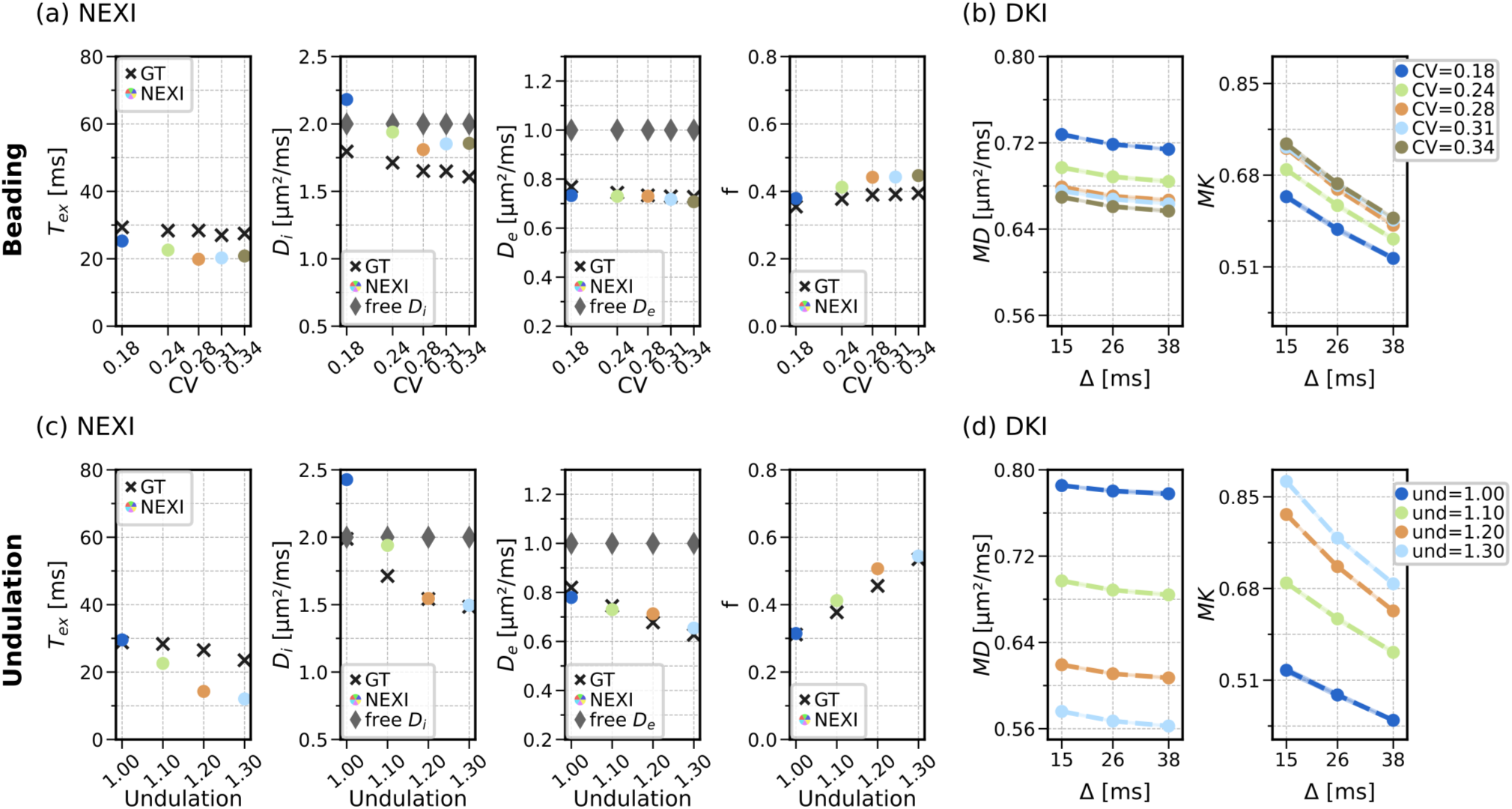
NEXI performance with neurite irregularities. NEXI parameters *t_ex_*and *f* showed limited sensitivity to neurite beading, showing only a small increase in bias (from -20% to -24% and from +9% to +13% respectively) as beading level (CV) rose from 0.24 to 0.34 (a). *D_e_* remained constant and *D_i_* in general decreased with increasing beading, consistent with restricted intracellular diffusion (a), which was similarly reflected in a decrease of MD and an increase of MK (b). Similar trends were observed for undulation, both in the NEXI (c) and DKI (d) estimates, although in this case *D_e_* decreased with increasing undulation. Black cross markers indicate ground-truth (GT) values and gray diamond markers indicate *D_i_* and *D_e_*, the intrinsic free diffusivities specified in the simulator. Beading level was defined as the coefficient of variation of the radius, given by 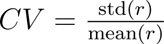 (H. H. Lee et al., 2019), and undulation was defined as the ratio between the total neurite length and its end-to-end distance.

NEXI estimates *t_ex_*and *f* showed limited sensitivity to beading, and although some bias was present, it increased only marginally with stronger beading (from -20% to -24% for *t_ex_* and from +9% to +13% for *f*) as beading (CV) increased from 0.24 to 0.34.

*D_e_*, which was consistently lower than the intrinsic diffusivity due to neurite packing, as noted previously, agreed with the ground-truth. *D_e_* remained relatively stable across all beading levels, with a slight decrease as beading increased, likely due to increased hindrance in the extracellular space. Neurite beading was correctly captured by the *D_i_* estimates, systematically decreasing as beading increased, consistent with reduced water mobility in the neurites caused by larger beads (Budde & Frank, 2010; H.-H. Lee et al., 2024; Nguyen-Duc, Brammerloh, et al., 2026). Some fluctuations in *D_i_* were observed at larger bead sizes, similar to *t_ex_*. This may be attributed to the slight increased orientation dispersion observed in these substrates with higher beading.

Consistent with these NEXI estimates trends, diffusion kurtosis metrics showed a decrease in MD and an increase in MK with increasing beading, reflecting the increased hindrance and non-Gaussian diffusion caused by the irregularities (Fig. 3b). The time dependence of both MD and MK remained consistent across all beading levels (Table 4a). Consequently, the corresponding exchange times were also similar across beading levels and were, on average, approximately 10 ms larger than the corresponding NEXI estimates (Table 5a).

**Table 4:** Slopes of MD and MK time dependence (in 10^3^ *µ*m^2^*/*ms^2^) for different neurite irregularities.

| (a) Beading (see Fig. 3b) |  |  | (b) Undulation (see Fig. 3d) |  |  |
| --- | --- | --- | --- | --- | --- |
|  | slope MD | slope MK |  | slope MD | slope MK |
| CV=0.18 | -0.59 | -4.97 | und=1.00 | -0.33 | -4.03 |
| CV=0.24 | -0.56 | -5.59 | und=1.10 | -0.56 | -5.59 |
| CV=0.28 | -0.53 | -6.18 | und=1.20 | -0.52 | -7.76 |
| CV=0.31 | -0.52 | -5.87 | und=1.30 | -0.58 | -8.24 |
| CV=0.34 | -0.56 | -5.97 |  |  |  |

**Table 5:** Exchange time *t_ex_* (ms): ground-truth (GT), NEXI-estimated, and MK time-dependence–derived values for different neurite irregularities.

| (a) Beading |  |  |  | (b) Undulation |  |  |  |
| --- | --- | --- | --- | --- | --- | --- | --- |
| | GT $t_{ex}$ | NEXI $\hat{t}_{ex}$ | MK $\hat{t}_{ex}$ | | GT $t_{ex}$ | NEXI $\hat{t}_{ex}$ | MK $\hat{t}_{ex}$ |
| CV=0.18 | 29 | 25 | 34 | und=1.00 | 29 | 30 | 35 |
| CV=0.24 | 28 | 23 | 32 | und=1.10 | 28 | 23 | 32 |
| CV=0.28 | 28 | 20 | 30 | und=1.20 | 27 | 14 | 26 |
| CV=0.31 | 27 | 20 | 33 | und=1.30 | 24 | 12 | 26 |
| CV=0.34 | 27 | 21 | 32 |  |  |  |  |

Unlike beading, increasing undulation reduced the accuracy of *t_ex_* (with bias increasing from -20% to - 49% as the undulation increased from 1.10 to 1.30). The estimates of *f* showed no systematic trend with increasing undulation. Note that because undulation altered the final volume fraction *f*, the increased bias in *t_ex_* may be partly attributable to changes in volume fraction rather than to undulation effects alone.

As in the case of beading, undulation increased hindrance to water diffusion within neurites, causing *D_i_* to decrease as undulation increased (Fig. 3c). In contrast to beading, undulation increased extracellular tortuosity, leading to a decrease in *D_e_* with stronger neurite undulations; which was accurately captured by the NEXI estimates of *D_e_*.

It is worth noting that in the cases of minimal beading and no undulation, *D_i_* estimates exceeded the intrinsic free diffusivity. This is likely due to the lower orientation dispersion in these two substrates, which is expected to introduce larger errors in diffusivity estimation (Pizzolato et al., 2023).

Undulation affected diffusion kurtosis metrics similarly to neurite beading (Fig. 3d): MD decreased, and MK increased with greater undulation, consistent with hindered diffusion within irregular neurites. Unlike beading, however, the time dependence of MD and MK increased slightly more with stronger undulation (Table 4b). The exchange times estimated from MK time dependence ranged from 26 ms at the highest level of undulation to 35 ms at the lowest undulation (Table 5b).

### 3.4 Impact of orientation dispersion and somas

NEXI assumes that neurites are randomly oriented; however, GM can exhibit varying levels of orientation dispersion (∼ 0.13–0.30) (Lampinen et al., 2020; Salo et al., 2021). Thus, we evaluated NEXI in a substrate with reduced orientation dispersion.

NEXI estimates *t_ex_*and *f* became less accurate with decreasing orientation dispersion (higher *p*_2_), with bias increasing from -20% to -39% for *t_ex_* and from +9% to +13% for *f* (Fig. 4a). These biases remained smaller than the uncertainty in *t_ex_* under realistic SNR conditions (Fig. 2c). *D_i_* remained stable (and, as before, lower than the intrinsic diffusivity due to neurite irregularities), whereas *D_e_* decreased with higher *p*_2_ as the extracellular space became more restricted perpendicular to the fibers, but remained accurately estimated (Fig. 4a).

**Figure 4:**
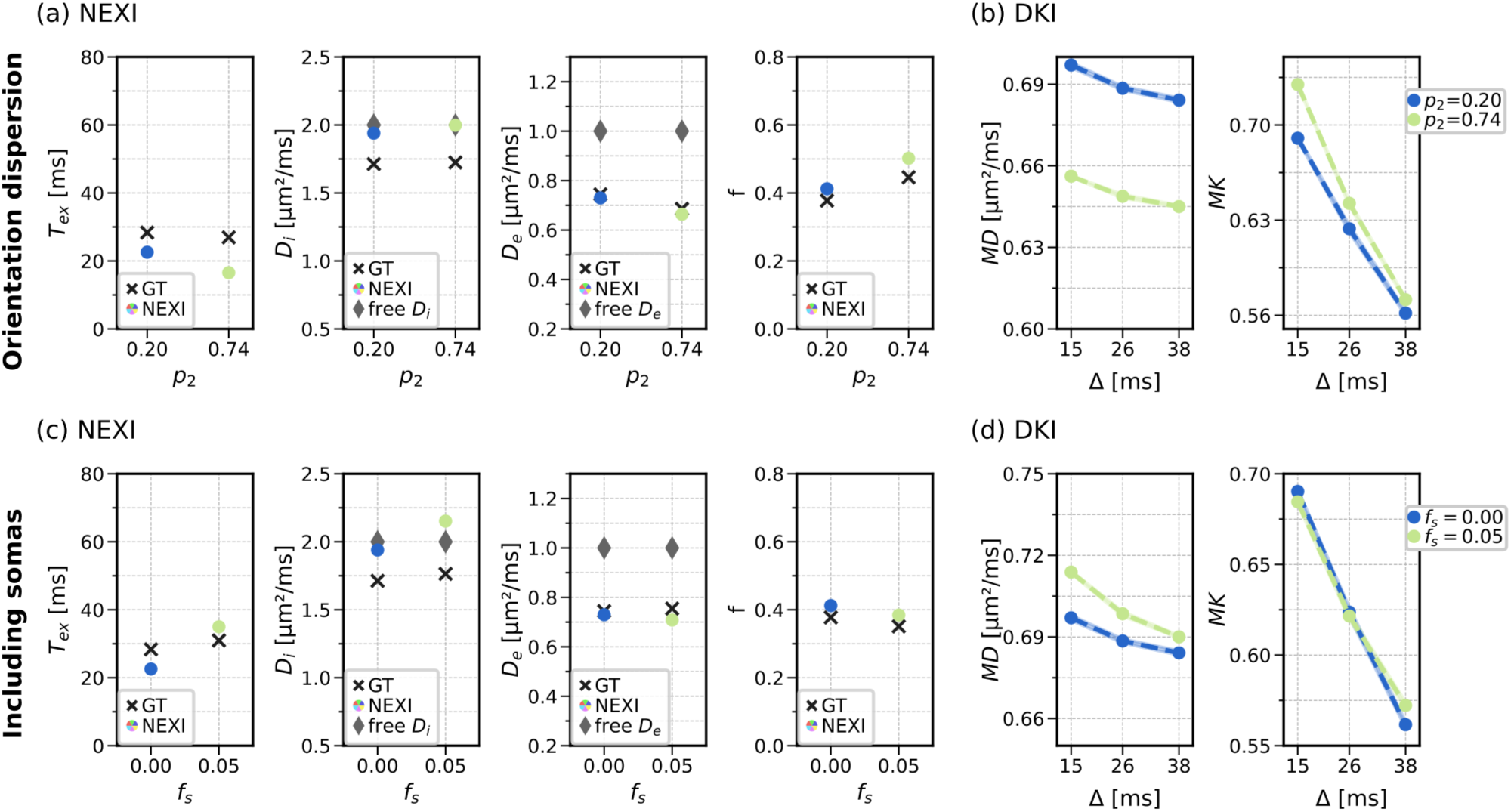
NEXI performance with different orientation dispersions and presence of somas. NEXI estimates *t_ex_* and *f* became less accurate with increased orientation dispersion (a). *D_i_* remained stable, while *D_e_* decreased with higher *p*_2_ as the extracellular space became more restricted perpendicular to fibers (a), lowering MD and slightly raising MK (b). The introduction of 5% somas, which are unaccounted for in NEXI, led to a similar magnitude bias in *t_ex_* and an overestimated *D_i_*, while *f* and *D_e_* were unaffected (c). MD increased due to the additional, less restricted diffusion within the somas, which was also reflected in a stronger MD time dependence. MK time dependence was unaffected (d). Black cross markers indicate ground-truth (GT) values and gray diamond markers indicate *D_i_* and *D_e_*, the intrinsic free diffusivities specified in the simulator.

Diffusion kurtosis metrics corroborated these findings: higher *p*_2_, corresponding to more aligned neurites, resulted in lower MD and slightly larger MK, reflecting reduced diffusion in the extracellular space (Fig. 4b). The time dependence of MK was more pronounced at low orientation dispersion (Table 6a), resulting in faster (this is, lower) exchange time estimates (Table 7a), consistent with the NEXI estimates.

**Table 6:** Slopes of MD and MK time dependence (in 10^3^ *µ*m^2^*/*ms^2^) for different orientation dispersions and presence of somas.

| (a) Orientation dispersion (see Fig. 4b) |  |  | (b) Presence of somas (see Fig. 4b) |  |  |
| --- | --- | --- | --- | --- | --- |
|  | slope MD | slope MK |  | slope MD | slope MK |
| $p_2=0.20$ | -0.56 | -5.59 | $f_s=0.00$ | -0.56 | -5.59 |
| $p_2=0.74$ | -0.48 | -6.86 | $f_s=0.05$ | -1.03 | -4.87 |

**Table 7:**
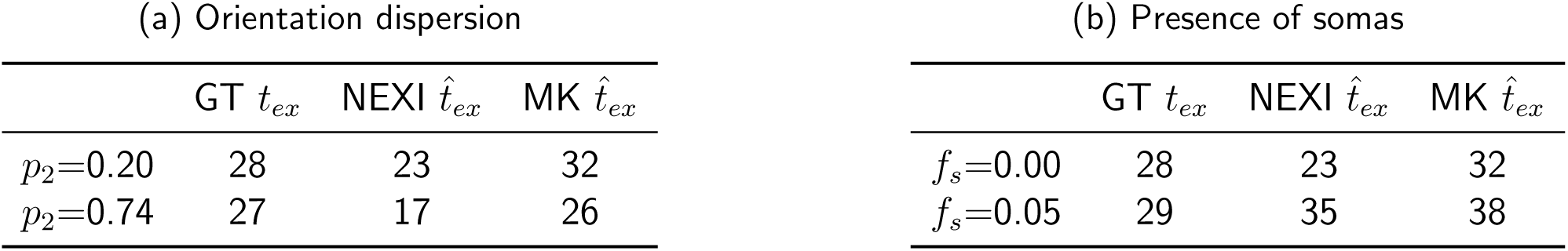
Exchange time *t_ex_* (ms): ground-truth (GT), NEXI-estimated, and MK time-dependence–derived values for different orientation dispersions and presence of somas.

Next, we evaluated the case of impermeable somas at a soma volume fraction of 10%, for which the signal was dominated by the somas. The synthetic diffusion MRI signal reflects a restriction-dominated regime with higher signal at longer diffusion times (Fig. S3a), which no longer reproduces the *in vivo* observation of an exchange-dominated decrease in signal with increasing diffusion time in GM (Jelescu et al., 2022; Olesen et al., 2022). As a result, the NEXI fit failed entirely. The signal time-dependence matched again experimental observations when permeability was also introduced in the soma membrane with the same value as that of neurites (Fig. S3a). In this case, an exchange time between intra- and extra-cellular spaces could be estimated, unlike when somas were impermeable. Nevertheless, the estimated exchange time *t_ex_* showed high standard deviation across repeats and low accuracy relative to the ground-truth (Fig. S3b). Indeed, a soma volume fraction of 10% in a substrate with neurite volume fraction of 30% represents an unrealistically high soma-to-neurite ratio of = 30%.

We therefore focused on the more realistic case of a soma volume fraction of 5% with soma–extracellular permeability *κ* = 20 *µ*m*/*s. Combined with a neurite volume fraction of 35%, this yields a soma-to-neurite ratio of 14%, which is more consistent with histological reports showing soma ∼ 9.4%, neurites ∼ 66%, and a corresponding soma-to-neurite ratio of ∼ 14% (Shapson-Coe et al., 2024). In this case, NEXI overestimated *t_ex_* by +21% (Fig. 4c), a bias comparable in magnitude, but opposite in direction, to that observed without somas (-20%). The introduction of somas led to an increase in *D_i_*. In contrast, *f* and *D_e_* remain largely unaffected by the presence of somas (Fig. 4c). Hence, NEXI estimates remain robust in the case of a realistic soma fraction.

Diffusion kurtosis metrics reflected the expected soma effects (Fig. 4d): the substrate containing 5% somas showed larger MD and exhibited increased MD time dependence due to highly restricted, non-Gaussian diffusion in the somas, as compared with the substrate without somas (Table 6b). MK exhibited the same exchange-dominated time dependence as in the case without somas, resulting in comparable exchange time estimates (Table 7b), which were higher when somas were present, similar to NEXI estimates.

### 3.5 Performance for clinical acquisition protocols

To reflect a realistic clinical acquisition, we simulated a protocol suitable for clinical use (Uhl, Pavan, Feiweier, et al., 2025) and evaluated SMEX instead of NEXI, as SMEX accounts for the non-zero duration of the diffusion gradients of this protocol. The performance with the clinical protocol (Fig. 5a) was overall similar to that observed with the preclinical protocol (Fig. 2a). However, the estimates of *t_ex_* and *D_i_* were less accurate with the clinical acquisition, with biases in *t_ex_* of -36%, -70%, and -20% for *κ* = 10, 20, and 30 *µ*m*/*s, respectively. A higher bias in *t_ex_* is observed at low permeabilities (in particular *κ* = 20 *µ*m*/*s) compared with Fig. 2a, which may be attributed to the shorter diffusion times used here (28.3 ms). Likewise, at *κ* = 20 *µ*m*/*s, both *f* and *D_e_* exhibited increased bias.

**Figure 5:**
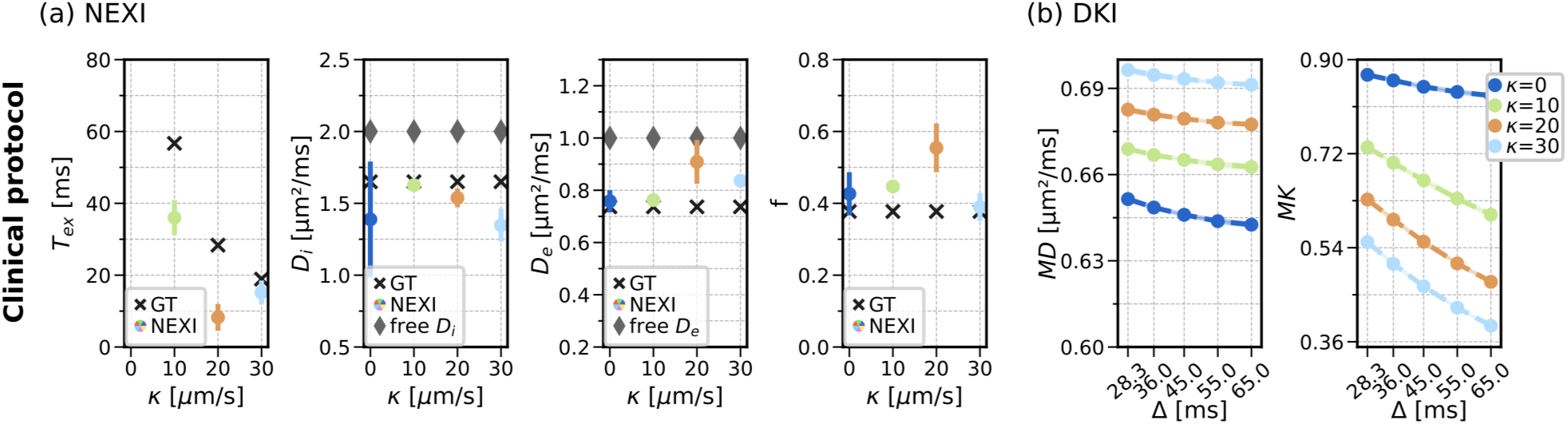
SMEX performance with a clinical protocol. SMEX parameter *D_e_*and *f* generally agreed with trajectory-derived values(black markers), with some deviations observed at *κ* = 20 *µ*m*/*s; both *D_i_* and *D_e_* were lower than intrinsic diffusivity (gray markers), reflecting hindered diffusion from packing and neurite irregularities (a). *t_ex_* followed theoretical predictions, with higher bias for *κ* = 20 *µ*m*/*s. MD showed weak time dependence (please note the limited range of the y-axis), while MK exhibited a pronounced time dependence that increased with increasing permeability (b), consistent with the preclinical protocol (Fig. 2b). Black cross markers indicate ground-truth (GT) values and gray diamond markers indicate *D_i_* and *D_e_*, the intrinsic free diffusivities specified in the simulator.

Similarly to the preclinical protocol, the diffusion kurtosis metrics also reflected the effect of exchange: MK showed a clear time dependence, whereas MD time-dependence was much weaker (Fig. 5b).

## 4 Discussion

In this work, we evaluated the performance of NEXI in GM-like substrates that incorporate realistic neurite features such as beading and undulation, with different orientation dispersions and inclusion of somas, across different levels of permeability. We observe that NEXI exchange time estimates mirror the different permeability levels well and remain largely accurate under realistic neurite irregularities, altered orientation dispersion, and the presence of somas, despite some bias. This study builds on the important challenge of validating biophysical models, ensuring that their assumptions and estimates remain reliable under realistic microstructural conditions.

### 4.1 Substrate generation

To conduct this validation, we generated GM-like numerical substrates with the CATERPillar toolbox (Nguyen-Duc, Brammerloh, et al., 2026). One of the geometric features of GM that we aimed to reproduce was high orientation dispersion, though this proved challenging with CATERPillar, as this toolbox was primarily designed for WM. To approximate different dispersion levels, we simulated two substrates by combining two or three fiber populations; however, intermediate dispersion levels could not be generated, limiting a full assessment of orientation dispersion impact on NEXI estimates.

Since somas are not accounted for in NEXI, we examined their potential influence on parameter estimates by including soma compartments in our simulated substrates. The soma-to-neurite ratio in our simulations (14%) was consistent with histological findings (∼ 14%) (Shapson-Coe et al., 2024). However, the absolute soma and neurite volume fractions we were able to achieve (5% and 35%, respectively) remained at the lower end of reported values (somas: 5–40%; neurites: 40–75%) (Motta et al., 2019; Shapson-Coe et al., 2024). As a result, the substrates contained a larger extracellular space than typically reported in *ex vivo* fixed tissue, which constitutes a limitation of our simulations. Nonetheless, it is noteworthy that the neurite fraction estimations of NEXI were reasonably accurate across all substrates and configurations, and that the ground-truth values achieved in our synthetic substrates match, in fact, the NEXI estimates of *f* in the GM *in vivo* (for rat and human) (Chan et al., 2025; Jelescu et al., 2022; Uhl, Pavan, Feiweier, et al., 2025). At the same time, previous *ex vivo* or *postmortem* measurements of *f* using NEXI or SMEX yielded much higher *f* estimates, compatible with fixed tissue histology or electron microscopy analyses (Hertanu et al., 2023; Olesen et al., 2022). It is therefore plausible that our synthetic substrates are well representative of *in vivo* GM tissue.

Overall, there is a need for improved numerical substrate generation frameworks capable of packing neurites more densely while allowing realistic tuning of GM cellular morphology (Aird-Rossiter et al., 2026; De Riedmatten et al., 2024; Villarreal-Haro et al., 2023).

Despite these limitations, we were able to generate substrates that approximate key aspects of GM microstructure: they exhibit high orientation dispersion, reasonably packed neurites, realistic neurite morphology through beading and undulation, tunable permeability and, in one case, inclusion of somas. Although not fully realistic, these substrates are sufficient for evaluating NEXI, as they capture the main relevant GM features.

### 4.2 NEXI performance in realistic GM-like substrates

Realistic neurites (with beading and undulation) introduced some time dependence in MD, which were more pronounced with stronger undulation (*ɛ* = 0.4, und=1.30) (Fig. 3b,d). This behavior suggests a contribution from *structural disorder* (Novikov et al., 2014) within this diffusion-time range (15–38 ms). For these diffusion times, the characteristic longitudinal diffusion length, 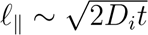, is on the order of 8–12 *µ*m. This length scale is much larger than the neurite radius (∼ 1 *µ*m), and is therefore insensitive to neurite radius. In contrast, it is comparable to the characteristic length scale of neurite undulations and beading (*ℓ* ∼ 10 *µ*m), as reported for substrates with comparable amplitude (*A* = 0.3) and undulation (*ɛ* = 0.4) (Nguyen-Duc, Brammerloh, et al., 2026). This indicates sensitivity to along-neurite structural variations within this diffusion-time range.

The presence of structural disorder (manifested as within-compartment diffusivity time dependence) violates a key assumption of the Kärger model (Gaussian diffusion in each compartment, thus diffusivity time independence per compartment) and could therefore challenge the validity of NEXI. Importantly, NEXI estimates remained consistent with the expectations even under such conditions (Fig. 3a,c). In fact, MK, whose time dependence has been shown to arise from competing mechanisms (structural disorder and exchange between compartments (Fieremans et al., 2010; H.-H. Lee et al., 2020; Novikov et al., 2014)), was strongly modulated by permeability (Fig. 2b,d) and less by neurite irregularities such as beading (Fig. 3b), but with large undulations also inducing a non-negligible MK time dependence (Fig. 3d). Overall, these results highlight the predominant influence of intercompartmental exchange in GM, despite neurite irregularities, underscoring the validity of the central model characteristic of NEXI (Fieremans et al., 2010; Jelescu et al., 2022).

#### Effect of permeability, beading and undulation

To further probe this relationship, we then examined the effect of membrane permeability in the most realistic default substrate. Overall, NEXI estimates remained robust over the examined range of permeabilities. In particular, the accuracy of extracellular diffusivity and exchange time remained fairly stable across permeability levels. However, NEXI estimates of intracellular diffusivity increased while estimates of neurite volume fraction decreased as permeability increased (Fig. 2a,c). This is likely because, at high permeability (when exchange across the membrane is larger), there are fewer intracellular molecules that have a unidirectional diffusion signature (they also experience extracellular space with 3D mobility). Thus there is seemingly a reduced proportion of water confined in unidirectional sticks but whose diffusivity is closer to the free diffusivity as they do not encounter any membrane restriction.

In terms of accuracy, the estimated exchange time also aligned with theoretical predictions based on the underlying substrate geometry, despite a negative bias that remained roughly constant across permeability levels (∼18%). Exchange times estimated from the MK time dependence also agreed well with the ground-truth (Figs. 2–3). At membrane permeability *κ* = 20 *µ*m*/*s, NEXI exchange time was underestimated by ∼20%, with this bias remaining stable across all beading levels. This suggests that NEXI may remain applicable in conditions with elevated neurite beading, such as cytotoxic edema and, in particular, ischemic stroke (Budde & Frank, 2010), while still allow reliable estimation of exchange times. Undulation, on the other hand, affected the accuracy of exchange time estimation more markedly. The heavier underestimation possibly stems from different neurite “segments” appearing to exchange, and thus decreasing the exchange time (similarly to an effect of spine vs neurite shaft exchange proposed in (Şimşek et al., 2025)). An additional contributing factor may be the increased neurite volume fraction associated with substrates exhibiting higher undulation.

#### Effect of orientation dispersion

A less dispersed neurite configuration (higher *p*_2_) also yielded a stronger underestimation of *t_ex_* (−20% to −39%), which could be due to the departure from the model assumptions of an isotropic extracellular space or a higher probability for extracellular molecules to encounter neurite membranes. Nevertheless, this bias remained smaller than the uncertainty observed under realistic SNR conditions.

Decreasing orientation dispersion also slightly increased the bias in *f* (+9% to +13%). Although these biases appear large, it is important to note that the corresponding *p*_2_ values are considerably higher than those typically reported for GM (Lampinen et al., 2020; Salo et al., 2021).

#### Effect of acquisition protocol

Remarkably, *t_ex_* was more heavily underestimated when using the clinical protocol of longer gradient pulses, longer diffusion times and lower b-values – though not consistently across permeability values. This is possibly counterintuitive, as the range of diffusion times is thought to act as a filter on the measurable exchange times: we thus would have expected an overestimation of the exchange time with the clinical protocol. The driver of this trend remains to be investigated.

#### Effect of somas

Finally, *t_ex_* was overestimated by ∼21% when 5% somas were included. The effect of somas on the estimation of *t_ex_* is not very pronounced in this range compared to uncertainties at realistic SNR (see below).

Beyond this effect on *t_ex_*, somas also influenced other parameters. The inclusion of somas further increased *D_i_*, which is surprising given that, based on the stable *f* estimate, NEXI seems to impart the soma compartment to the extracellular water, as previously suggested (Jelescu et al., 2022). This overestimation of *D_i_* could explain, in part, the high estimates found *in vivo* for this parameter (sometimes close to the free water diffusivity of 3.0 *µm*^2^*/ms*). These estimates are reduced back to more biologically plausible values when somas are included in the model such as in SANDIX (Jallais et al., 2025; Mezzano et al., 2025). Taken together, these findings indicate that, in this diffusion time regime, NEXI yields robust parameter estimates despite not explicitly accounting for somas.

It is important to note that the simulated voxel used soma properties consistent with values obtained from microscopy-reconstructed neurons (gamma distribution for the soma radius with a 7.1 *µ*m mean and 3.6 *µ*m standard deviation (Aird-Rossiter et al., 2026)). However, these results are based on the limited histological data available, and regional as well as inter-species variability in soma size and volume fraction may lead to different NEXI estimates. Indeed, when the soma fraction was increased to 10%, the behavior of the synthetic signals changed quite substantially. At a given b-value, the MR signal was higher with longer diffusion times, which is a clear signature of restriction dominating over exchange. As a result, the NEXI fit for *t_ex_* failed, with the estimate hitting the upper bound. When these 10% soma were assigned the same permeability as the neurite membrane – which is not implausible biologically – the simulated MR signals retrieved their expected behavior of exchange dominating over restriction (lower signal for longer diffusion times) and an estimate for *t_ex_* was output, though overestimated at 111 ms.

This prompts a few considerations. Firstly, while 10% soma does not seem excessive in absolute terms, here this substrate had an artificially high soma-to-neurite ratio because the neurite fraction could not be increased proportionally beyond 30%. This explains why, in the case of these soma being impermeable, the signal signature changed so radically and departed also from experimental observations: in practice, exchange was always observed to dominate over disorder *in vivo* in GM (i.e. lower signal for longer diffusion times). Secondly, this also points to the fact that soma membranes are likely also permeable, and a proper GM model should account for neurites, soma, but also for their permeable membranes (and not just for permeable neurites) and possibly for the exchange soma-neurite. Our results suggest that permeable soma have less of a restrictive signature on the signal and could explain over-estimations of soma density or size in models of impermeable spheres such as SANDI (Palombo et al., 2020).

Finally, it is noteworthy that, in most substrates and configurations, *D_i_* was quite systematically overestimated by NEXI, ranging close to the free intracellular diffusivity value (2.0 *µm*^2^*/ms* in our simulation) but larger by ∼ 12% than the ground-truth value obtained from the trajectories. The trajectory-based estimate is lower than the free diffusivity value as it accounts for the neurite irregularities. This is again consistent with the fact that *in vivo* estimates for *D_i_* are overall quite high (Jallais et al., 2025; Jelescu et al., 2022; Uhl, Pavan, Feiweier, et al., 2025), and should perhaps be interpreted through the prism of this known over-estimation.

### 4.3 Noise

To evaluate NEXI’s robustness to noise, we assessed its behavior with and without added noise across different SNR levels. In the presence of moderate noise (SNR = 50, typically exceeded in preclinical acquisitions and within the range commonly observed in GM in clinical acquisitions) both the variability and bias of NEXI estimates increased (Fig. 2c). The increased variability in *t_ex_* at lower permeabilities reflects the limited range of diffusion times used (15–38 ms). Compared to the noise-free case, the overall trends of parameter values were nevertheless preserved. At lower SNR (SNR = 20, Fig. S2) the variability and bias increased substantially. These results suggest that NEXI remains reliable at moderate SNR levels, which are realistic for cortical SNR in clinical acquisitions, but becomes less reliable as SNR decreases. Overall, we demonstrated the robustness of NEXI under realistic noise conditions.

Similarly, the diffusion kurtosis metrics MD and MK exhibited not only substantial variability but also a bias compared to the noise-free case, with MD systematically higher under noisy conditions. The inaccuracy of MK estimators has been reported previously under conditions of low SNR or limited sampling directions (Veraart et al., 2011, 2013).

In the noiseless case, MK exhibited a marked dependence on diffusion time, whereas MD showed only a weak trend. At SNR=50, the time dependence of MD became highly variable but MK slopes remained statistically different from zero and their trend across permeabilities was preserved. At SNR=20 though, the time dependence of both MD and MK became unstable and departed from expected physical behavior due to noise.

Such noisy time dependence, particularly for MD and also observed *in vivo* (H.-H. Lee et al., 2020), may explain the difficulty in identifying clear and consistent time-dependent trends in experimental GM data, and may contribute to the discrepant findings across studies. Indeed, some reports show little MD time dependence (Jelescu et al., 2022; H.-H. Lee et al., 2020), typically interpreted as a lack of structural disorder and, together with marked MK dependence, as evidence of inter-compartmental exchange as the dominant mechanism in GM. Others, however, report a non-negligible MD time dependence, which violates the Kärger model assumptions, implying that exchange times cannot be reliably estimated under such conditions (Mougel et al., 2024). Our study shows that realistic levels of neurite irregularities, in the form of varying degrees of beading and undulation, do introduce diffusion time-dependence. However, this time-dependence due to structural disorder is weak enough to be obscured by noise, and, even in the noiseless case, did not interfere with acceptable and meaningful estimates of NEXI parameters. Indeed, NEXI retained sensitivity to permeability, neurite fraction, and effective intra-neurite diffusivity.

### 4.4 Future perspectives

In this work, the exchange effects simulated reflected only membrane-mediated exchange between the intra- and extracellular compartments; however, other exchange mechanisms exist. As discussed above, dendritic spines can give rise to exchange-like diffusion signatures, but their contribution is expected to be limited under the diffusion-time regimes considered here (Chakwizira et al., 2025; Şimşek et al., 2025). Nevertheless, our work showed that undulation may cause neurites to behave as different exchanging “segments” or “domains” and contribute to lowering the estimated exchange time below the membrane permeability value on timescales more comparable to preclinical and clinical dMRI acquisitions than the spine-shaft exchange.

Other potential exchange effects include water exchange between somas and neurites, and between connected neurite branches (similar to undulations) (Ianus et al., 2021). These mechanisms could influence the interpretation of NEXI parameters in GM, as the estimated exchange time may reflect a combination of permeative and geometric exchange processes.

Therefore, future work should incorporate additional GM microstructural features, including spines and other forms of intracellular exchange.

As alluded to before, another important direction for future work is the use of numerical substrates that better reproduce GM, allowing for denser neuronal packing, and enabling more realistic assessments of model validity under realistic conditions.

## 5 Conclusions

Here, we evaluated the NEXI model with numerical substrates that mimic GM microstructure. These substrates were generated with realistic neurite features, such as beading and undulation, varying degrees of orientation dispersion, and the inclusion of somas, across different membrane permeability levels. We showed that NEXI reliably estimates the exchange time across permeability levels and successfully disentangles exchange from other microstructural effects. These findings support the model’s potential to capture exchange in GM. Future work should aim to further improve substrate realism to better represent the complexity of neuronal morphology. Overall, we show that NEXI is suitable for *in vivo* microstructure mapping in the healthy and diseased human brain.

## Data and Code Availability

NEXI code is publicly available at https://github.com/Mic-map/graymatter swissknife, and the CATER-Pillar code is available at https://github.com/Mic-map/CATERPillar. Synthetic data are available upon request.

## Author Contributions

**Rita Oliveira:** Conceptualization, Methodology, Formal analysis, Investigation, Writing – Original Draft, Writing – Review & Editing. **Jasmine Nguyen-Duc:** Conceptualization, Methodology, Software, Writing – Review & Editing. **Malte Brammerloh:** Conceptualization, Software, Writing – Review & Editing. **Ileana Jelescu:** Conceptualization, Writing – Review & Editing, Funding acquisition.

## Supplementary Material

**Figure S1:**
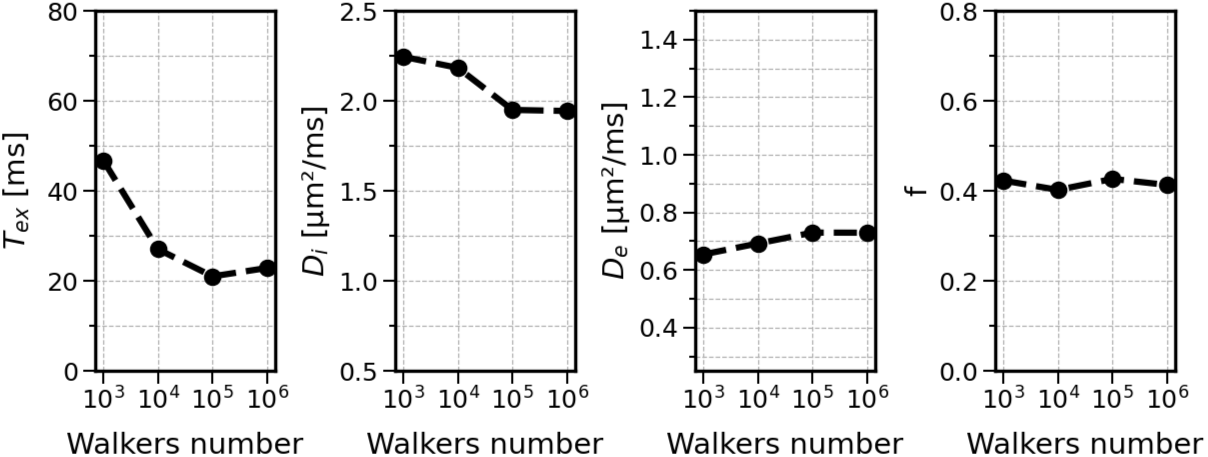
NEXI estimates as a function of the number of walkers simulated. Increasing the number of walkers beyond 10^5^ did not substantially improve the estimates, as most parameters reached a plateau.

**Figure S2:**
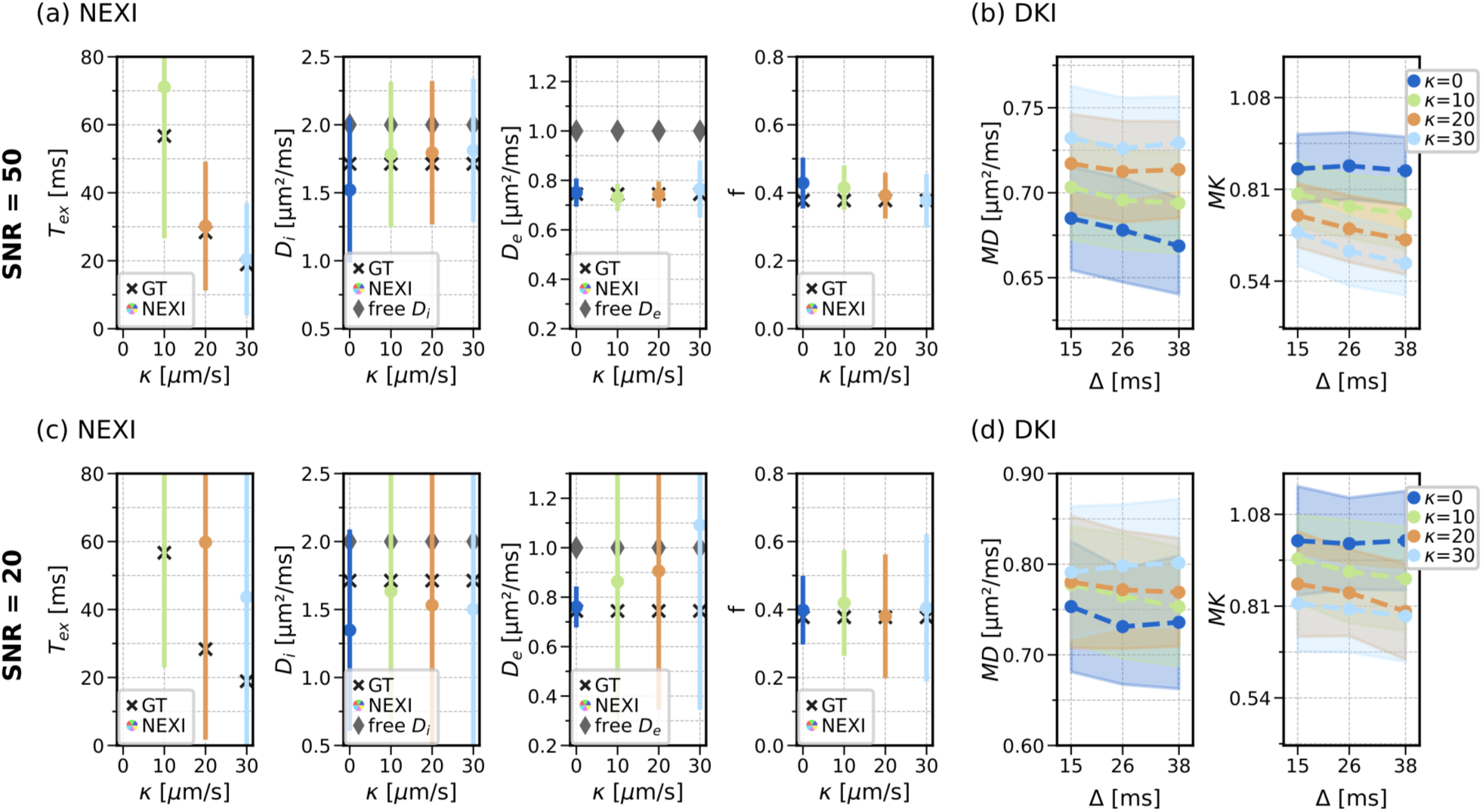
NEXI performance across permeability levels with SNR = 50 and 20. NEXI (a, c) and DKI (b, d) estimates with added noise at SNR = 50 and SNR = 20, respectively. With decreasing SNR, NEXI estimates show higher variability and increased bias. The time dependence of MD and MK becomes unstable and departs from expected physical behavior at lower SNR levels. Black cross markers indicate ground-truth (GT) values and gray diamond markers indicate *D_i_*and *D_e_*, the intrinsic free diffusivities specified in the simulator.

**Figure S3:**
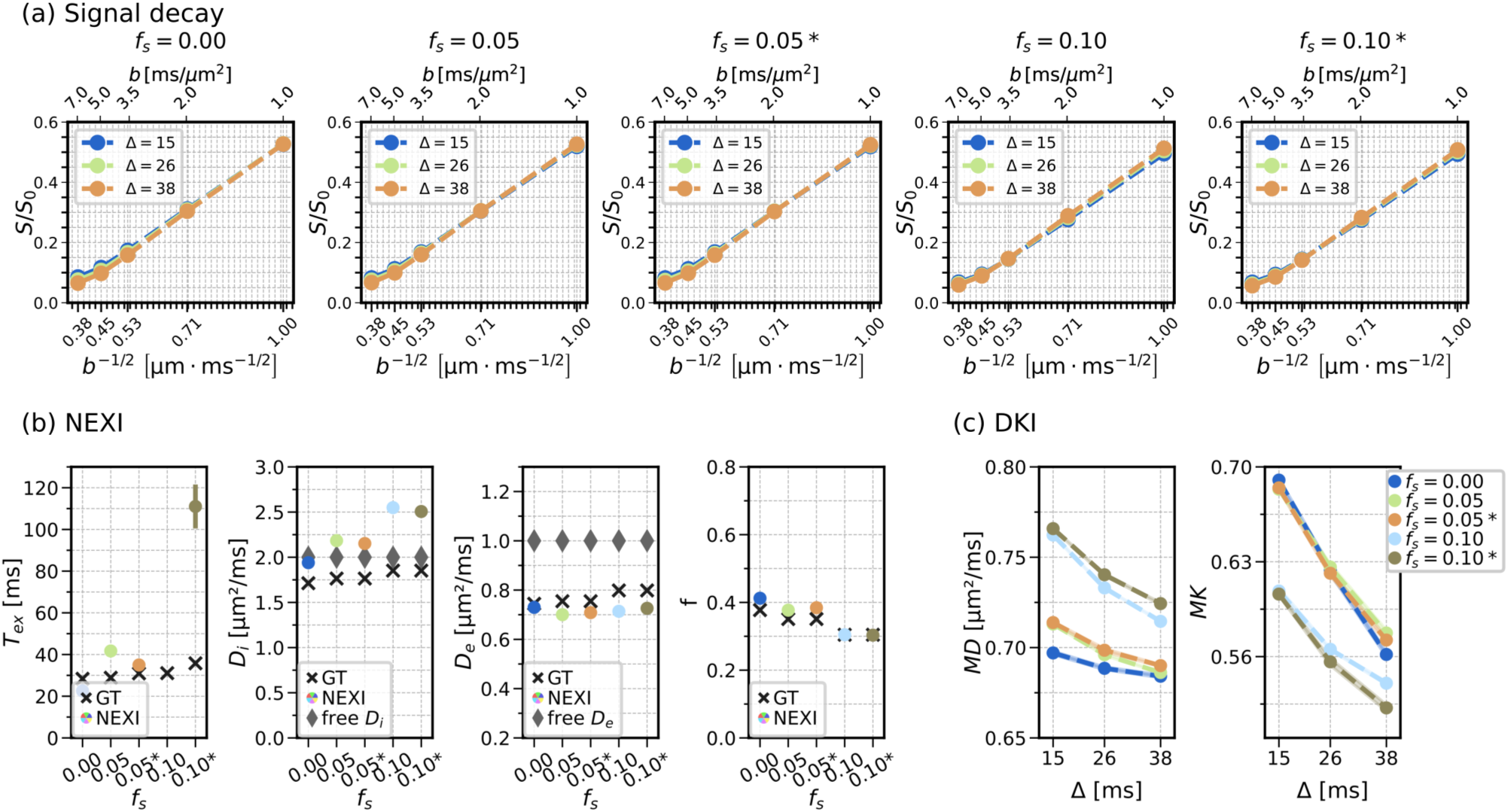
NEXI performance in the presence of 5% and 10% somas. We evaluated the following conditions: *f_s_* = 0; *f_s_* = 0.05 impermeable somas; *f_s_* = 0.05 with soma–extracellular exchange at 20 *µ*m*/*s; *f_s_* = 0.10 impermeable somas; and *f_s_* = 0.10 with soma–extracellular exchange at 20 *µ*m*/*s. The asterisk (*) indicates cases where soma–extracellular exchange is included, whereas the remaining cases assume impermeable somas. All cases used neurite membrane permeability of 20 *µ*m*/*s. Signal decays for these conditions are shown in (a). For the case of impermeable somas, as the soma fraction increased, the soma contribution progressively dominated the overall signal, and the diffusion MRI signal no longer reproduced the *in vivo* observation of decreasing signal with increasing diffusion time (Jelescu et al., 2022; Olesen et al., 2022), preventing estimation of an exchange time (b). The *in vivo*–like behavior was recovered when soma–extracellular exchange was introduced, although the resulting *t_ex_* estimates exhibited substantial variability and bias (b). The DKI metrics showed similar time dependence in the exchange and no-exchange cases, although MD increased and MK decreased for higher soma volume fractions due to the greater contribution of less restricted spherical diffusion (c). Black cross markers indicate ground-truth (GT) values and gray diamond markers indicate *D_i_* and *D_e_*, the intrinsic free diffusivities specified in the simulator.

